# Anisotropic stretch biases the self-organization of actin fibers in multicellular Hydra aggregates

**DOI:** 10.1101/2024.10.02.616220

**Authors:** Anaïs Bailles, Giulia Serafini, Heino Andreas, Christoph Zechner, Carl Modes, Pavel Tomancak

## Abstract

During development, groups of cells generate shape by coordinating their mechanical properties through an interplay of self-organization and pre-patterning. Hydra displays a striking planar pattern of actin fibers at the organism scale, and mechanics influence the morphogenesis of biological structures during its pre-patterned regeneration. However, how mechanics participate in the formation of an ordered pattern from a totally disordered state remains unknown. To study this, we used cellular aggregates formed from dissociated Hydra cells, which initially lose all actin polarity yet regenerate a long-range actin pattern. We showed quantitatively that the actin meshwork evolves from a disordered symmetric state to an ordered state in which rotational symmetry is broken, and translation symmetry is partially broken, with the nematic and smectic order parameters increasing over days. During the first hours, the actin meshwork displayed spatial heterogeneity in the nematic order parameter, and ordered domains separated by lines of defects progressively grew and fused. This suggests that local cell-cell interactions drive the transition from disorder to order. To understand the mechanism of ordering, we perturbed the tissue’s physical constraints. We showed that while topology and geometry do not have a direct effect, anisotropic stretch biases the emerging orientation of the actin meshwork within hours. Surprisingly, although a Wnt protein gradient is expected to play a role in the actin ordering, the stretch-associated alignment happened without a Wnt enrichment. This demonstrates the role of tissue mechanics in the alignment of the actin fibers during the disorder-to-order transition.

## Introduction

While self-organization plays an important role in the development and regeneration of animals, it is often coupled with some level of pre-patterning or external cues, such as the deposition of maternal RNAs or sperm entry. *In vitro* cellular systems such as organoids have shown that, in principle, self-organization is sufficient to create cellular patterns (Clevers, 2016; Yang and Liberali, 2021). However, mechanical forces can influence pattern formation in development (Collinet and Lecuit, 2021; Shyer et al., 2017), and the mechanisms by which cells use mechanics to self-organize in the absence of pre-patterning and external stimuli remain unknown. A vintage “organoid”, *Hydra vulgaris*, can form a complete animal from an aggregate of 10,000 to 100,000 dissociated cells within days and without cell proliferation or external inputs (Gierer et al., 1972; Noda, 1970). Its main body axis is patterned by a gradient of Wnt protein activity and is characterized by a striking actin fibers alignment (**Fig. 1A**). This raises the possibility that the actin meshwork interacts with biochemical patterning during body axis regeneration (Braun and Keren, 2018), making it an ideal model system to study the mechanochemical principles of cellular self-organization. Recent biophysical studies in Hydra have focused on the dynamics of topological defects in the actin meshwork and its effect on morphogenesis using regeneration of tissue fragments (Maroudas-Sacks et al., 2024a, 2024b, 2021; Ravichandran et al., 2024) that inherit the actin pattern from the adult axis (Maroudas-Sacks et al., 2024a; Shani-Zerbib et al., 2022). Here, we leveraged Hydra aggregates made from dissociated cells to erase the initial pattern, enabling the study of the transition from a disordered to an ordered actin meshwork from cells to organism scale.

**Fig. 1.**
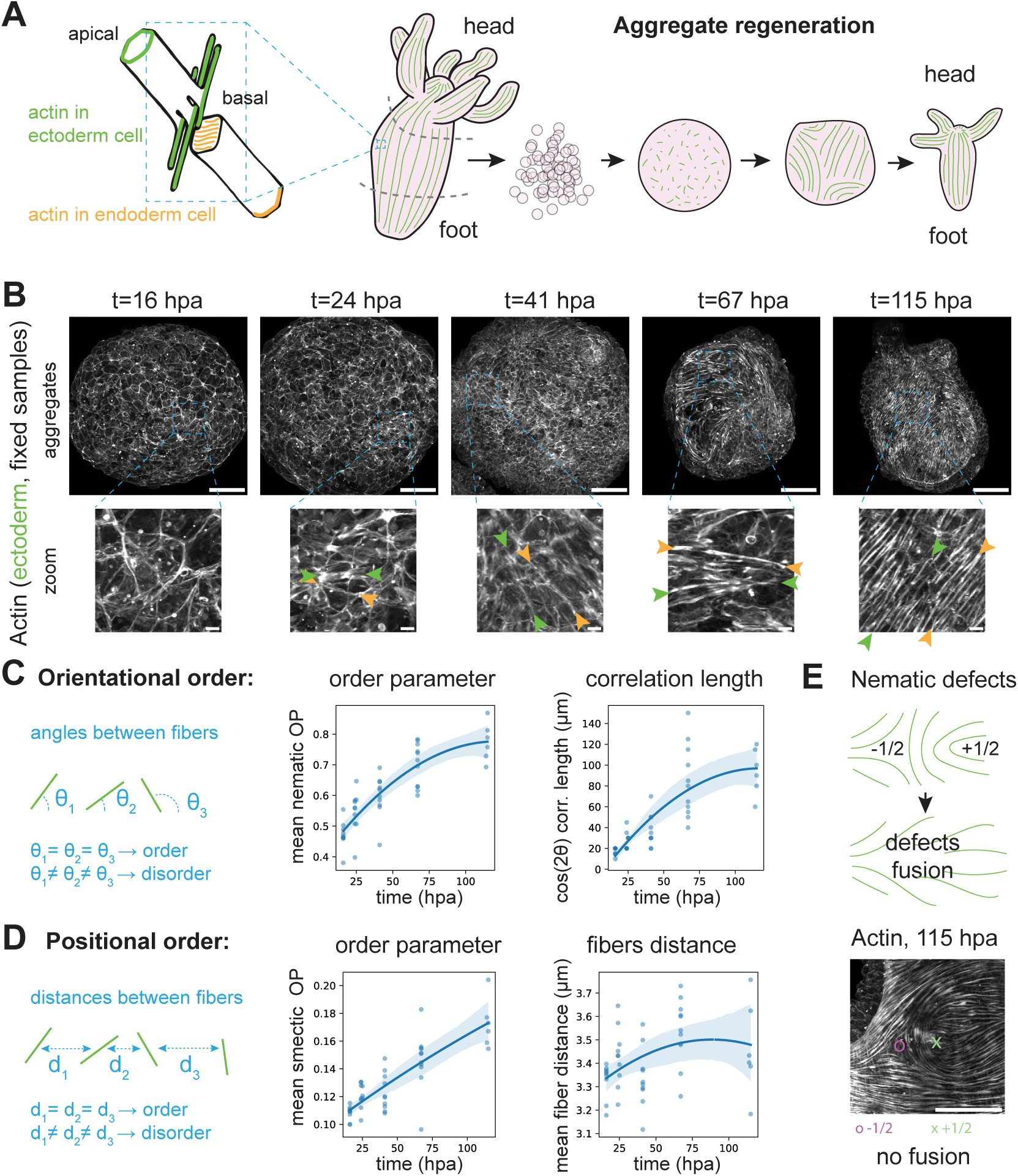
Actin fibers orientational and positional order increase with time. (**A**) Left, schematic of actin fibers within an ectodermal and endodermal cell, adapted from (*10*). Right, schematic of actin fibers during the process of aggregate regeneration. (**B**) Confocal microscopy images of actin in the ectoderm (LifeAct-GFP) of 5 fixed aggregates at 5 time points in aggregate regeneration. Hpa: hours post aggregation. Bottom: zoomed-in images. The green and orange pair of arrows point to the tips of a single actin fiber. Fibers at later time points are longer and more parallel. (**C**) Left, 1D schematic of the concept of orientational order, which measures whether the angles θ between neighboring fibers are similar: the more the angles are similar, the more ordered the fibers are. Center: orientational order parameter (nematic OP) with time. Right: correlation length with time, defined as the distance with a 20% correlation between cos(2θ) values. (**D**) Left: 1D schematic of the concept of positional order, which measures if the distances d between neighboring fibers are similar: the more the distances are similar, the more ordered the fibers are. Center: positional order parameter (smectic OP) with time. Right: characteristic fibers distance. For the 4 plots in (C) and (D), one dot is the spatial average of one aggregate, with n=46 aggregates from N=3 batches, and the blue line is a 2^nd^-order polynomial regression as a visual guide, with a lighter area representing the confidence intervals. (**E**) Confocal microscopy zoomed-in image of actin (phalloidin) in a 115 hpa aggregate, showing an example of a pair of −1/2 (purple dot) and +1/2 (green cross) defects. n=8 WT aggregates from N=1 batch fixed at 115 hpa displayed 5 pairs of similar defects in total. All microscopy images are maximum-intensity stack projections. Scalebars are 100 microns, except in zoom views, where they are 10 microns.

The adult Hydra has a simple body plan consisting of the oral-aboral axis, with a head surrounded by tentacles at one end and an adhesive foot at the other (Bode, 2011; Galliot, 2012; Vogg et al., 2019). The oral-aboral axis and the head are patterned by genes from the canonical Wnt pathway (Broun et al., 2005; Gee et al., 2010; Hobmayer et al., 2000). The body column of Hydra is made of two layers of epitheliomuscular stem cells, which give Hydra its regenerative abilities. On the contrary, the head and foot cells are terminally differentiated and cannot regenerate. Throughout the body plan, the ectodermal layer outside and the endodermal layer inside are separated by an extracellular matrix (ECM) consisting mainly of collagen. On the basal side, facing the ECM, both the ectoderm and the endoderm cells have contractile actin fibers (myonemes), which allow Hydra to move by contracting its body column (**Fig. 1A**). In the ectodermal layer, actin fibers are aligned with the oral-aboral axis, while the endodermal ones are orthogonal. The actin fibers in different germ layers also differ by size, with ectoderm fibers extending beyond the cell diameter, while the endoderm ones are thinner and shorter (Aufschnaiter et al., 2017; Seybold et al., 2016).

Aggregates can be made from dissociated cells of the body column, thus lacking the biochemical organizer activities of the head and foot. During the first phase, which lasts around ten hours, endodermal and ectodermal cells spontaneously sort and reestablish the epithelial bilayer and the ECM (Cochet-Escartin et al., 2017; Skokan et al., 2020). A lumen then appears at the center of the sphere, which undergoes cycles of inflations followed by rupture of the epithelium and abrupt deflation (Fütterer et al., 2003; Kücken et al., 2008; Mombach et al., 2001; Sato-Maeda and Tashiro, 1999). After a few days, protrusions and tentacles appear (**Movie S1**), followed by the formation of one or several functional heads. The resulting multi-headed Hydra composed of mixtures of cells from many individuals can feed and reproduce, re-establishing organismal identity. The basal actin fibers are lost following the dissociation step (Seybold et al., 2016). Therefore, unlike in Hydra tissue fragments, the actin alignment directional cues are erased in the aggregates. The tissue-scale fiber pattern is later established *de novo* (Seybold et al., 2016); however, the dynamics and biophysical mechanisms of this process remain unknown.

In this work, we leverage the Hydra aggregates devoid of biochemical and mechanical pre-patterns to study how the actin fibers pattern evolves from a disordered to an ordered phase over time, using a combination of imaging, image analysis, and perturbations. We describe the physical parameters governing the transition from disorder to order, investigate the influence of physical constraints on this transition, and demonstrate a direct effect of anisotropic stretch on actin alignment emergence.

## Results

### Actin fibers’ positional and orientational order increase with time

Using cells expressing LifeAct-GFP in the ectoderm (Aufschnaiter et al., 2017) to label actin, Seybold et al. observed that actin fibers are entirely lost during cell aggregate formation, form again at 12 hours post aggregation (hpa), align at the cell scale by 36 hpa, and start to align at the tissue scale at 36-48 hpa (Seybold et al., 2016) (**Fig. 1A**). We build on this initial qualitative description by imaging aggregates fixed at different time points after aggregation across 5 days (at 16, 24, 41, 67, and 115 hpa) and quantifying actin fiber alignment (**Fig. 1B**). The fibers were apparent at the cell scale from 24 hpa and acquired an alignment at the tissue scale from 41 hpa. This alignment improved with time (**Fig. 1B**). We developed an image analysis pipeline to extract relevant quantitative descriptors of the degree of order of the fibers (see Methods and **Fig. S1A**). First, we were interested in the loss of orientational symmetry among the fibers. When the fibers point in random directions, the actin field is isotropic or rotationally symmetric. This symmetry is broken when some directions are favored, initially at the scale of a patch of cells, then at the scale of the entire aggregate. This phenomenon is measured by the nematic order parameter *S* =< cos(2*θ*) > (see Methods), which is 0 for a fully disordered actin pattern and 1 for a fully ordered one. In Hydra aggregates, the nematic order parameter mean increases steadily from 0.47±0.05 at 16 hpa to 0.77±0.06 at 115 hpa (samples mean±std, **Fig. 1C left**, **Fig. S1B**). The order parameter value at 16 hpa was not 0, even though very few basal actin fibers are present at this stage because the junctional apical actin contributes to the order parameter. The scale at which the alignment persists can be measured by the characteristic length extracted from the autocorrelation function of the fibers’ angles (see Methods). It increases from 17.2±3.6 microns to 94±22 microns over the duration of the experiment (**Fig. 1C right**, **Fig. S1B**). This increase does not follow the same trend as other measures based on the intensity of the actin meshwork (**Fig. S1C**). Hence, we provided a quantitative estimate of the global increase in the actin nematic order of Hydra aggregates and showed that this increase occurs progressively over five days. The transition from disorder to order appeared smooth, without any noticeable abrupt change.

In addition, we observed that while fibers do not initially have a fixed distance between them, their position, and hence the distance between them, become more regular with time. In other words, the actin field is initially translational symmetric, and this second symmetry may also be broken later when the fibers acquire an array-like pattern along one dimension. This is analogous to the layers formed in the smectic phase of liquid crystals, which have a fixed distance between them. More precisely, the Hydra actin fibers on a 2D surface are analogous to the 2D layers in a 3D liquid crystal volume. Therefore, we devised a second (smectic) order parameter associated with the characteristic distance between parallel fibers (Methods, **Fig. S1A**), which, similarly to the nematic order parameter, spans values between 0 and 1. This smectic order parameter increased steadily as the aggregate matured, although with much smaller values than the nematic order parameter, from 0.108±0.005 to 0.172±0.018 (**Fig. 1D**). The associated averaged characteristic distance between fibers (or smectic-like layers) remained almost constant at 3.42±0.15 microns (all time samples mean±std). Importantly, such a smectic-like organization of the actin fibers in tissues has consequences for the expected evolution of the nematic defects. For example, −1/2 and +1/2 nematic defects close to one another in a nematic field are expected to be attracted and fuse, hence disappearing. However, in a smectic field, the presence of layers between the defects can prevent their fusion. Indeed, we observed examples of −1/2 and +1/2 nematic defects close to one another in late WT aggregates when the head and tentacles are already formed (**Fig. 1E**). We would expect such defects to have resolved, as they do not correspond to biological structures. Looking closely at the fibers, some are seamlessly continuous from one cell to the next at late stages (**Fig. 1B**, zoom of 115 hpa). We speculate that the fibers of adjacent cells are progressively attached by adhesion complexes at the level of desmosome-like junctions (Seybold et al., 2016), which leads to a solid supracellular fiber (or layer) that cannot be easily disassembled.

Taken together, we show quantitatively that the maturation of aggregated Hydra cells into a functional organism involves a transition of actin fibers orientation and, to a lesser degree, actin fibers position to a higher-order state. Due to the formation of supracellular actin fibers, this ordered state is more akin to a smectic crystal than a nematic one at late stages.

### Spatiotemporal dynamics of the nematic order increase

In order to resolve the dynamics of actin fibers order increase with a few minutes temporal resolution, we then turned to light-sheet microscopy. We imaged living free-floating and agarose-confined LifeAct-GFP aggregates during the 20 hours (from around 26 hpa to 46 hpa) when the actin fibers evolved from cell-scale order to tissue-scale order (**Fig. 2A**, **Fig. S2A**, **Movie S2**). During the imaging period, the aggregates underwent one to three inflation cycles (**Fig. S2B**). Outside these global events, we observed some local, seemingly reversible, constrictions of patches of cells (**Movie S2**). When we quantified the nematic order, the mean of the orientational order parameter increased over one period of inflation, corroborating the observations on fixed data (**Fig. 2C**). The angle autocorrelation length also increased, regardless of whether it is measured in microns or as the percentage of the aggregate diameter (**Fig. 2D,E**). Aggregates confined by agarose had, on average, a similar nematic order evolution (**Fig. S2B-F**). The inter-aggregate variability observed in fixed samples can thus be partly attributed to samples being fixed at different moments in the inflation-deflation cycles, technical variability, and aggregate size variability (**Fig. S2B**).

**Fig. 2.**
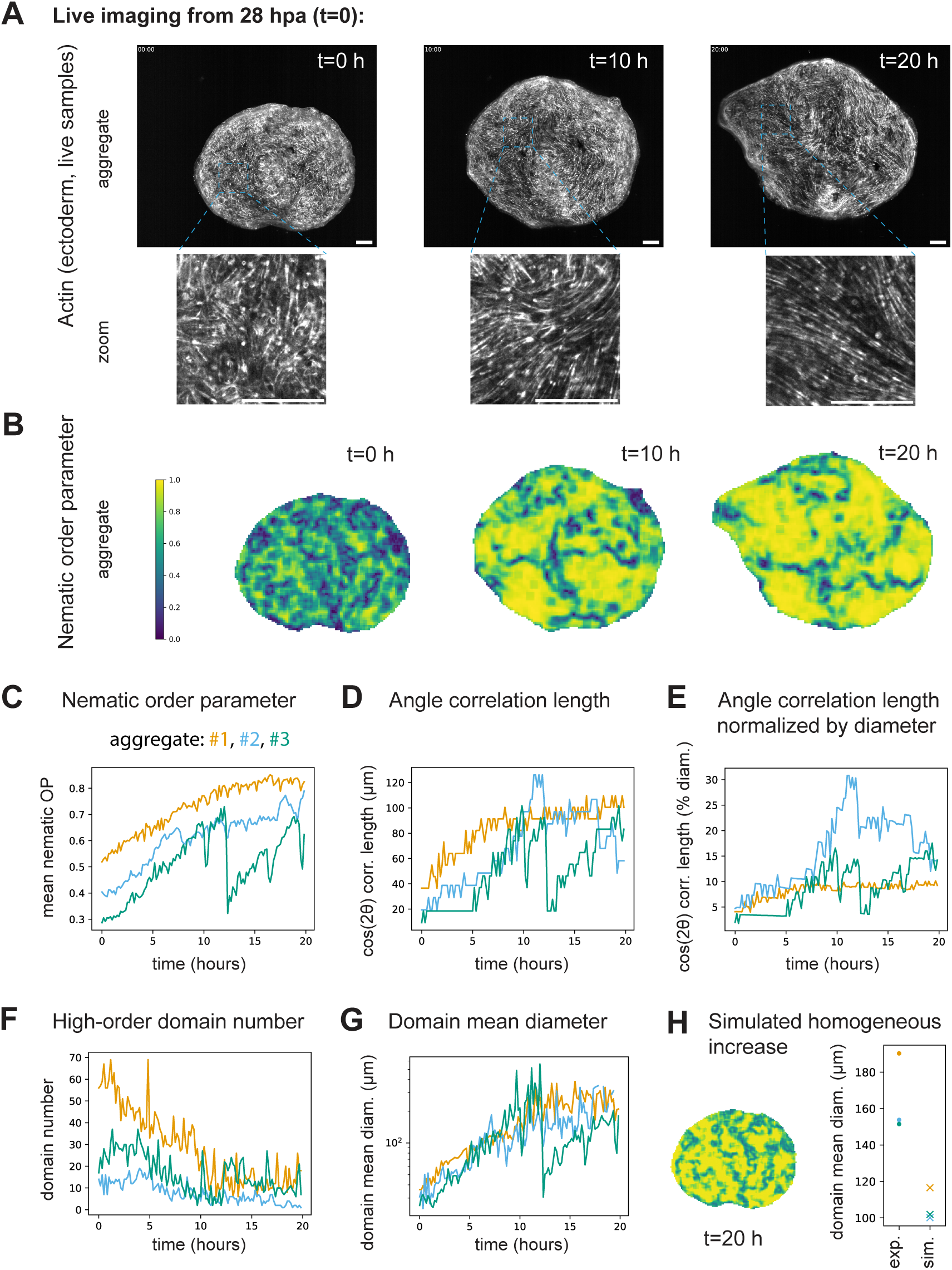
Spatiotemporal dynamics of the nematic order increase. (**A**) Three time points of a light-sheet microscopy movie of actin in the ectoderm (LifeAct-GFP) of a free-floating aggregate starting at 28 hpa (hours post aggregation). Bottom: zoomed-in images. (**B**) Corresponding spatial measurements of the orientational order parameter (nematic OP). Left: color bar, color code from 0 to 1). (**C**) Spatial average of the nematic order parameter with time of n=3 live LifeAct-GFP aggregates in water from N=3 batches. (**D**) Correlation length (defined as in Fig. 1C) with time of the same samples. (**E**) Correlation length normalized by the sample’s diameter at each time point. (**F**) High-order domains (defined as in **Fig. S2G**) number with time for the same samples. (**G**) Mean high-order domain diameter with time of the same samples. (**H**) Left: inferred order parameter spatial distribution after 20h of homogeneous increase starting from the spatial distribution at t=0h of the aggregate in (A). Right: comparison of the mean diameters of high-order domains between real aggregates and their simulated homogeneous growth counterpart. All microscopy images are maximum-intensity stack projections. All scalebars are 100 microns.

Interestingly, analysis of the order parameter derived from live imaging of the aggregates in water revealed spatial heterogeneities that evolve with time (**Fig. 2B**). We defined high-order as the areas of the aggregates with an order parameter above a threshold (**Fig. S2G**, Methods) and analyzed the changes of the connected high-order domains over time. The number of such connected domains decreased with time while their area increased (**Fig. 2F,G**). Domain fusion events were observed and detected in the distributions of individual domain areas (**Fig. S2J**). Eventually, the aggregates comprised only a few ordered domains separated by disordered stripes, also called lines of defects (**Fig. 2B**). Such lines of defects could not appear if a concentration gradient aligned the actin fibers. In addition, the domains had higher growth rates and were bigger than they would have been if the order parameter had increased homogeneously (**Fig. 2H**). This suggested that the transition from disordered to ordered nematic field did not occur uniformly across the aggregate but instead by growth and fusion of higher-order domains, likely driven by cell-cell local interactions.

The spatial heterogeneities may be initiated or amplified by differences in epithelial tissue state (e.g., epithelial junctions and ECM maturity) and actin fibers maturity differences. However, the order parameter did not correlate locally with the actin fibers’ meshwork intensity (**Fig. S2K**), which is expected to be proportional to actin density. We thus proceeded to identify what mechanical or physical parameters could facilitate or bias the ordering.

### Topology and geometry perturbations of the actin meshwork

As the alignment of the ectoderm actin meshwork in Hydra fragments is characterized by the striking formation of a +1 defect correlating with the head organizer (Maroudas-Sacks et al., 2021), we sought to perturb the *topology* of the tissue in the aggregates to see if it has an impact on the nematic field. At first, we created an extrinsic hole by laser puncture of the tissue of ∼20 hpa aggregates (when the tissue scale order is not in place yet) and imaged them until the actin meshwork reached a large-scale nematic order. Upon puncture, the aggregate deflated, and rapid wound healing occurred. After that, the normal inflation-deflation cycle restarted. However, the actin meshwork did not develop defects, not even transient, at the position of the laser puncture (**Fig. S3A, Movie S4**). This shows that a temporary tissue topological defect such as a hole does not directly affect the basal actin fiber alignment. To sustain the topological perturbation, we pierced 16 hpa old aggregates with a thread and left the aggregates to develop with the thread inside. This created two permanent topological defects in the tissue (**Fig. 3A, Fig. S3B**). Previous work used thin wires to generate tissue topological defects in Hydra fragments, with no measurable bias in the position of the head (Livshits et al., 2017). However, fragments conserve an intrinsic tissue planar polarity and alignment of actin, which could be stable to this perturbation. Hence, we repeated the experiment with aggregates and 50 microns diameter nylon threads (i.e., of the area of a patch of ∼3-4 cells). Following the placement of the thread, at two days post aggregation (dpa), actin (stained with phalloidin) was still partly disorganized (**Fig. 3A**). Later, at 4 dpa, the actin pattern was well-defined and long-ranged, and ectoderm fibers were easily distinguishable from endoderm ones by their diameter. At this stage, we manually measured the angle of the ectoderm actin pattern with respect to the thread, in proximity to both holes and on both sides of the samples (**Fig. 3A**). The results showed a significant difference to a uniform distribution (Kolmogorov-Smirnov ks-test, p=0.0074), and parallel alignment was measured in ∼50% more cases compared to orthogonal alignment (**Fig. 3B, Fig. S3C**). This indicated that the thread was introducing a bias in the actin meshwork. However, this could be caused by the side effects of the piercing process, such as wounding or deformation of the tissue. To uncouple these effects from topology, we used the unique property of the aggregates: by introducing the thread at 0 hpa, when the cells have not built adhesion and the tissue is not formed, topological defects can be created by displacing cells without damaging a tissue (**Fig. 3C**). In this case, we observed a smaller and non-significant bias of actin alignment with respect to the thread axis (ks-test p=0.071, ∼25% difference between parallel and orthogonal alignment) (**Fig. 3D, Fig. S3C**). Therefore, persistent defects associated with tissue wounding or deformation do slightly bias the actin pattern, while tissue topological defects alone do not significantly bias the actin orientation.

**Fig. 3.**
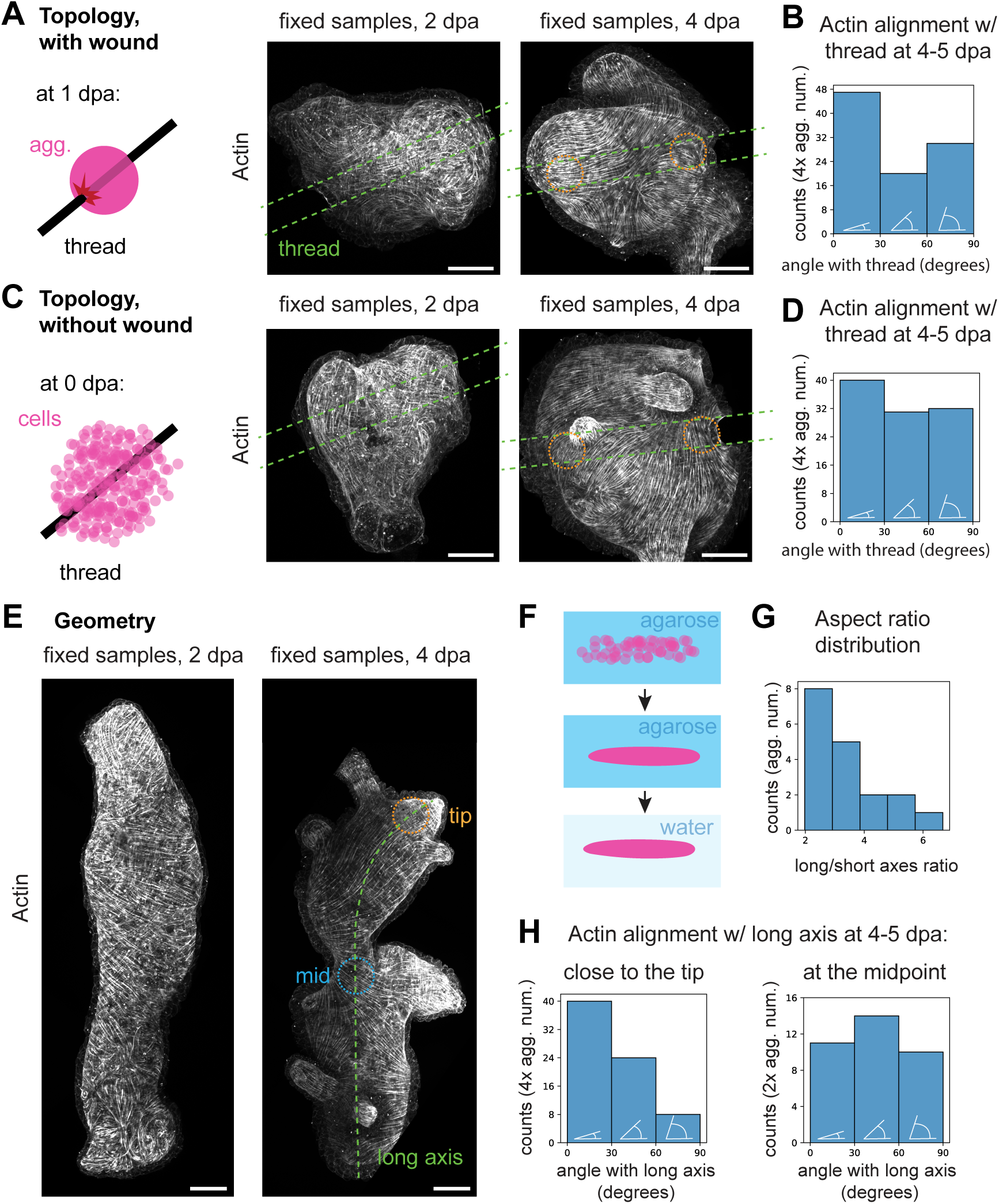
Topology and geometry perturbations of the actin meshwork. (**A**) Left, schematic of the topology perturbation experiment, with tissue wounding. Right, microscopy image of actin (phalloidin) in such a perturbed aggregate fixed at 2 dpa (days post aggregation), and another at 4 dpa. The thread contours are the green dashed line drawn from the bright field channel (displayed in **Fig. S3**). The orange dashed circles show the approximative regions where the manual measurements are performed. (**B**) Manual measurements histogram of the actin angle with respect to the thread, in the regions close to the thread as highlighted in (A), at 4-5 dpa, binned in parallel (0-30°), diagonal (30-60°), and orthogonal (60-90°) categories, represented with white schematics. 97 measures from n=25 aggregates, N=3 batches. (**C**) Left, schematic of the topology perturbation experiment, without tissue wounding. Right, microscopy image of actin (phalloidin) in such a perturbed aggregate fixed at 2 dpa and 4 dpa. The thread contours are the green dashed line drawn from the bright-field channel (displayed in **Fig. S3**). The orange dashed circles show the approximative regions where the manual measurements are performed. (**D**) Manual measurements histogram of the actin angle with respect to the thread, in the regions close to the thread as highlighted in (C), at 4-5 dpa, binned in 0-30°, 30-60°, and 60-90° categories. 103 measures from n=27 aggregates, N=3 batches. (**E**) Microscopy image of actin (phalloidin) in a perturbed aggregate fixed at 2 dpa and at 4 dpa. Green dashed line: longest curvilinear axis. Orange dashed circle: tip region measured. Blue dashed line: midpoint region measured. (**F**) Schematic of the geometry perturbation. (**G**) Histogram of the aspect ratios of the samples analyzed. (**H**) Manual measurements histogram of the actin angle with respect to the long axis, in the regions at the tip (left plot) and midpoint (right plot) as highlighted in (E), at 4-5 dpa, binned in 0-30°, 30-60°, and 60-90° categories. 72 measures (for the tips) or 32 measures (for the midpoint) from n=18 aggregates, N=3 batches. All microscopy images are maximum-intensity stack projections. All scalebars are 100 microns.

Another possible source of bias that could orient the actin domains is the overall geometry of the aggregate. Geometry has been shown to orient actin in mammalian epithelial cells cultured on cylinders with radii <40 microns (Yevick et al., 2015) and up to 200 microns (Bade et al., 2017, p. 20). To test the effect of *geometry* on actin alignment in Hydra, we created aggregates with a large aspect ratio by centrifugating cells in a 200-micron diameter capillary. We then placed the elongated aggregates (“sausages”) in 2% agarose overnight to prevent their rounding up (**Fig. 3F**). Once the epithelialization of the aggregate was complete and the cells did not sort significantly anymore (>16 hpa) (Cochet-Escartin et al., 2017), we removed the aggregates from their agarose mold. The high aspect ratio persisted over at least 4 days (**Fig. 3E,G**), resulting in aggregates in which the geometry has been changed. This geometry was now the new preferred state of the tissue and was likely not associated with residual stress. The “sausage” aggregates had a small curvature (a large radius) along the long axis and a higher curvature (a shorter radius) along the short axis (**Fig. S3D**). At 2 dpa, the actin meshwork did not seem to align with the long or short axis, and the fibers were still not aligned at a large scale (**Fig. 3E**). At 4 dpa, the tips of the elongated aggregates displayed a significant bias in ectoderm actin orientation compared to a uniform distribution, with five times more parallel alignment with the long axis compared to the orthogonal one (ks-test p=2.1E-5) (**Fig. 3E,H, Fig. S3D**). However, the midpoints did not show such a bias (**Fig. 3H, Fig. S3D**), and we observed frequent orthogonal rather than parallel alignment of actin at the narrower part of the tissue (**Fig. S3F**). This showed that the anisotropy in curvature cannot account for the bias in favor of the parallel orientation at the tip, as the difference between the principal curvature values is much higher at the midpoints than at the tip. Having excluded curvature, we analyzed the distribution of +1 defects as proxies for the head organizer position across the “sausage” aggregate to understand why actin aligns with the tips. We measured the average half distance between two +1 defects to obtain the average random distance between a point and the closest defect. Next, we measured the distance between tips and the closest +1 defects. Since the result shows that the +1 defects are significantly closer to the tips than other points in the aggregate (Mann-Whitney U-test p=0.0021) (**Fig. S3G**), the bias in actin orientation measured at the tip reflects the head-organizer attraction to the tip rather than actin orientation in response to curvature.

### Stretch aligns the actin meshwork independently of Wnt3

What could attract the head organizer to the tip of the “sausage” aggregate? We hypothesized that the organizer position is biased due to the tip region experiencing higher stretches during inflation. To test if deformation affected actin, we proceeded to perturb *stretch* (here, meaning extensile strain) in the tissue specifically. The stretch in a control aggregate is approximately isotropic when the lumen inflates. To create an anisotropic deformation, we placed one-day-old LifeAct-GFP aggregates in agarose except for a 200-micron diameter cylindrical hole and imaged them live (**Fig. 4A**). During the aggregate inflation, the agarose restricted the stretch to the position of the hole and made it anisotropic (**Movie S5**, **Fig. 4B-D**). This dramatically affected the actin alignment, with fibers forming almost exclusively in the direction of the hole at the protrusion ‘neck’ (**Fig. 4F**). The effect extended to the aggregate center, where the actin fibers were mainly parallel or diagonal to the hole (ks-test p=0.013) (**Fig. 4E**). This alignment of the actin fibers was relatively fast, visible as early as 3.1 hours, and within 5.3 ± 3.5 hours (median±std, n=11, N=4) after agarose hole formation (**Movie S5**). The effect persisted over days at the ‘neck’ of the subset of aggregates, showing a lasting protrusion (ks-test p=1.7e-8) (**Fig. 4F**), but the alignment close to the center vanished progressively (ks-test p=0.082 at 2 dpa and p=0.33 at 4 dpa) (**Fig. 4E**). Since the aggregates were often seen rotating within the agarose confinement after 2 to 3 days (**Movie S6**), repositioning the anisotropic stress, the orientation of actin alignment didn’t persist in some cases. However, it was sometimes possible to observe striking and stable actin alignment in the protrusion within the agarose hole, even at 4 to 6 dpa with or without the presence of a head (**Fig. S4A**). Hence, this experiment suggested that the *stretch* of the tissue has a much more profound impact on the actin pattern than *topology* or *geometry*.

**Fig. 4.**
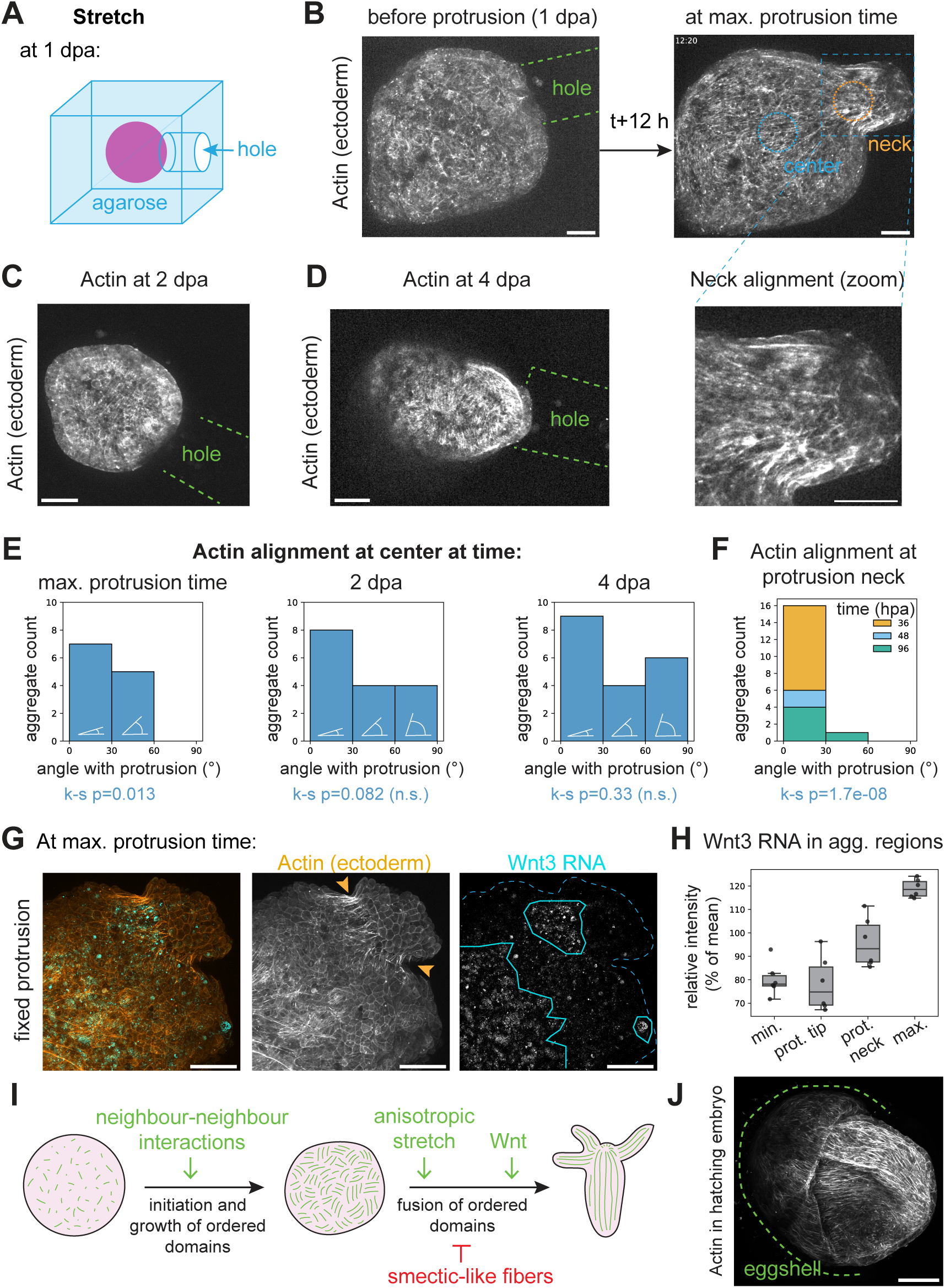
Stretch aligns the actin meshwork. (**A**) Schematic of the stretch perturbation experiment. (**B**) Spinning disk (SD) microscopy images showing actin in the ectoderm (LifeActGFP) in such a perturbed aggregate, before the protrusion (just after the creation of the hole) and after 12h (around the maximum protrusion time). Green dashed line: hole contours from bright-field images in **Fig. S4B**. Blue circle: example of the center measurement region. Orange circle: example of the ‘neck’ measurement region. Bottom right: zoomed-in image of the neck. (**C**) SD microscopy live image of actin in the ectoderm (LifeActGFP) in a 2 dpa (days after aggregation) aggregate placed in the agarose hole at 1 dpa. (**D**) SD microscopy live image of actin in the ectoderm (LifeActGFP) in a 4 dpa aggregate placed in the agarose hole at 1 dpa. (**E**) Manual measurements histogram of the actin angle with respect to the center-hole axis, in the center region as highlighted in (B), binned in 0-30°, 30-60°, and 60-90° categories. Left: at maximum protrusion time, n=12 aggregates, N=4 batches. Center: at 2 dpa, n=16 aggregates, N=4 batches. Right: at 4 dpa, n=19 aggregates, N=4 batches. (**F**) Manual measurements histogram of the actin angle with respect to the protrusion axis, in the neck region as highlighted in (B). The measures of the aggregates showing a protrusion at three different time points are stacked. n=17 protrusions necks, N=4 batches. (**G**) HCR RNA *in situ* hybridization with a *Wnt3* antisense probe in two examples of aggregate protrusions fixed 6 hours after 1 dpa hole creation. Left: composite image. Center: ectodermal actin (LifeActGFP). The orange arrows show the neck sides. Right: background-subtracted Wnt3 channel. The cyan lines delimit three tissue areas strongly expressing Wnt3 signal. (**H**) Normalized Wnt3 background-subtracted intensity measurements in different regions: the minimal intensity region, the protrusion tip, the protrusion neck, and the maximal intensity region, in n=6 aggregates from N=2 batches. (**I**) Proposed model for actin alignment in aggregates. (**J**) Actin (phalloidin) in a hatching Hydra egg. The dashed green line shows the eggshell as per the bright-field image. All microscopy images are maximum-intensity stack projections. All scalebars are 100 microns.

We next asked whether the actin alignment induced by mechanical stretch also depends on chemical positional cues developing within the regenerating aggregate. In Hydra, it has been established that Wnt signaling aligns actin, as grafting a piece of head on the body column locally changes actin orientation (Wang et al., 2020). The *Wnt3* expression is increased by osmotically-induced stretch (Ferenc et al., 2021) and wounding (Cazet et al., 2021) and is present early in aggregates (Hobmayer et al., 2000). To investigate if the actin alignment upon protrusion formation is due to mechanically-induced *Wnt* expression, we imaged LifeAct-GFP aggregates in the local stretch-inducing confinement in agarose with a hole (**Fig. S4E**). We fixed them once a protrusion was prominent (i.e., 5-10 hours after agarose hole formation) and visualized *Wnt3* expression in these anisotropically stretched aggregates by whole-mount HCR RNA *in situ* hybridization with a *Wnt3* antisense probe. This probe reliably labels the head organizer in adults (**Fig. S4H**). Even after the HCR protocol, the protrusions were still visible, as well as the stretch experienced by the cells (**Fig. 4G**, **Fig. S4D,E,F**). We found that the protrusion tip had a similar *Wnt3* expression as the regions of the aggregate with the lowest *Wnt3* intensity. Although the ‘neck’ showed higher expression, it was not higher than in other parts of the aggregate (**Fig. 4H**, **Fig. S4F,G**). Hence, increased *Wnt* expression at the protrusion cannot explain the aggregate-scale actin alignment upon induced anisotropic stress.

Taken together, this data supports the role of mechanical cues, such as anisotropic stretch, in aligning actin during the self-organized symmetry-breaking process of regenerating Hydra aggregates.

## Discussion

Overall, we quantified the ordering of Hydra aggregates actin fibers on both long and short timescales. Our measurements revealed that orientational order increases smoothly over a long timescale. Although the positional order rises only slightly, it is associated with the formation of stable connected supracellular actin fibers, making it smectic-like (**Fig. 1**). On shorter time scales, we showed the presence of spatial heterogeneities in nematic order (**Fig. 2**). We then physically perturbed the aggregates to gain insight into the possible mechanisms of actin fiber alignment. Perturbations of topology and geometry suggested that both have an indirect effect (possibly through wounding or stretch) rather than directly affecting actin alignment (**Fig. 3**). Conversely, stretch had a strong and fast effect on aligning actin independently of Wnt3 (**Fig. 4**).

We thus propose that actin fibers self-organize in Hydra aggregates in two steps, possibly overlapping in time (**Fig. 4I**). First, at early stages, the actin fibers interact locally at the cell scale, and these neighbor-neighbor interactions lead to the rotational symmetry breaking in domains that grow over time. Second, aggregate inflation generates a stretch that helps domain fusion and eventually leads to the robust long-range ordering of the fibers at the tissue scale. At late stages, the actin large-scale orientation is modified by chemical signaling (in particular, Wnt) to generate +1 defects (otherwise unstable) matching the tissue genetic patterns (heads, feet, tentacles) and to globally align actin with the body axis on the organismal scale.

The exact nature of the local interactions of intracellular actin fibers leading to their alignment has yet to be discovered. It could be steric (with the mere fibers’ presence restricting the orientation of its neighbors), mechanical (with adhesion between fibers or to the ECM), or chemical (for example, planar cell polarity (PCP) signaling between neighboring cells). We provided evidence that symmetry breaking does not occur homogeneously but by growth and fusion of ordered domains (**Fig. 2**), likely amplified by cellular heterogeneities inherent to the aggregate preparation mode. Although the aggregate osmotic inflation provides approximately isotropic stretch, the heterogeneities in local actin order may make some regions more stretch-compliant and lead to localized anisotropic stretch. Since we showed that anisotropic stretch orients the actin fibers robustly and in a long-range (**Fig. 4**), this could lead to stable tissue scale actin orientation in the system. The smectic-like behavior of the actin fibers (**Fig. 1**) means that the actin meshwork is more ordered than a nematic phase and that another type of defect, dislocation defects, must be considered. The smectic-like fibers are likely to prevent the resolution of the lines of defects between domains. Hence, the role of stretch could be to overcome the associated energy barriers and promote the fusion of domains. The geometry does not influence the actin orientation locally (**Fig. 3**) but could modify the stretch pattern and the subsequent gene expression (e.g., *Wnt3*) through mechanosensing. The *Wnt3* pattern may emerge in the tissue early on, but the extent to which it crosstalks with actin remains unknown. At late stages, the topology inevitably constrains the global number of defects of the actin as a nematic field (with a total charge of +2 on a sphere). When more than two +1 defects are present, this leads to the formation of −1/2 defects, which are not driven by any known chemical pattern or mechanism. However, the topology would not have any strong local effect (**Fig. 3**).

Interestingly, the development of aggregates confined by agarose with a hole introduced on one side has some similarities with the process of Hydra egg hatching (Martin et al., 1997). In the latter case, the embryo, which consists of 2 layers of cells, is constrained by the eggshell until it breaks free from it on one side. A part of the embryo then inflates and stretches outside the eggshell before any morphological signs of a head with tentacles are visible. However, the head of the newly hatched Hydra does develop on the side that broke free from the eggshell (Martin et al., 1997). This suggests the intriguing possibility that aggregate regeneration in the agarose confinement recapitulates mechanisms used during embryonic development. Although hatching Hydra embryos are notoriously difficult to capture, we managed to catch and fix one specimen of Hydra embryo in the act of breaking free from the eggshell. Phalloidin staining revealed that actin is indeed aligned parallel to the protrusion in a hatching Hydra egg stretching half outside of its eggshell (**Fig. 4I**, **Fig. S4I**), while no +1 defects or other typical Wnt-related structures were present. Since Hydras never experience dissociation and reaggregation as part of their normal life cycle, the remarkable reformation capacity of the system may be due to artificially arriving at a state that resembles a physiological embryonic process. An exciting hypothesis is that the anisotropic stretch-induced actin alignment we uncovered participates in the embryo body axis formation or ensures its robustness.

Regardless of the developmental coincidences that make Hydra resilient to dissociation, the ordering of actin fibers in aggregates arises from a complex interplay of chemical, mechanical, geometrical, and topological self-organized constraints. The respective temporal order and causal role of chemical, mechanical, or topological cues are still unclear in Hydra tissue fragments (Maroudas-Sacks et al., 2024b; Ravichandran et al., 2024; Shani-Zerbib et al., 2022; Wang et al., 2020). Wnt signaling is a conserved mechanosensing pathway (Pukhlyakova et al., 2018) and could be at the origin of the mechanical effects on head formation observed in fragments, as experimental manipulations indicate (Ferenc et al., 2021; Maroudas-Sacks et al., 2024b; Weevers et al., 2024). Theoretical work also emphasized the importance of the coupling between nematic alignment and concentration gradient (Wang et al., 2023) or the role of stretch-based mechanical feedback into Wnt signaling (Maroudas-Sacks et al., 2024b; Mercker et al., 2015; Soriano et al., 2009; Weevers et al., 2024). Aggregates allow the disentangling of these respective contributions by monitoring the *de novo* emergence of both the Wnt concentration field and the actin field. We showed that actin pattern formation can be uncoupled from the Wnt pattern and provided evidence for a mechanism where the *anisotropy* in stretch instructs the emergence of actin order and axial alignment without a corresponding Wnt pattern. In the future, it will be important to assess if this mechanism has long-lasting consequences on the self-organization of the head organizer in the aggregates.

Beyond Hydra, cells grown in *in vitro* monolayers align their long axes akin to nematic liquid crystals (Elsdale and Wasoff, 1976), and nematic topological defects have been associated with tissue rupture or cell extrusion (Guillamat et al., 2022; Sonam et al., 2023). The molecular aspects of the interplay between actin fibers orientation, tissue anisotropic stretch or stress, and gene patterning are investigated in systems with powerful molecular toolkits. The cytoskeleton and associated proteins at focal adhesions are inherently mechanosensitive (del Rio et al., 2009; Greenberg et al., 2016; Han and Rooij, 2016; Harris et al., 2018), and actin fibers formed, modified, or oriented by stress are present in other systems: ventral stress fibers in mammalian migrating cells (Burridge and Guilluy, 2016; Livne and Geiger, 2016), actin bundles in the development of mammalian striated muscles (Fenix et al., 2018; Mao et al., 2022) or smooth muscles (Huycke et al., 2019), and apical stress fibers in insect epithelial tissues (López-Gay et al., 2020). It will be interesting to investigate if what we uncovered also extends to these systems, particularly if they form a smectic phase that prevents the fusion of aligned domains. Overall, we described the physical parameters governing the transition from disorder to order in a mechanochemical self-organization process and investigated the influences of physical constraints on this transition. While mechanochemical symmetry breaking also occurs *in vitro*, Hydra aggregate cellular polarity and actin fibers self-organize across scales from heterogeneous mixtures of cells to entire functioning and reproducing organisms, allowing to connect *in vitro* findings with developmental principles.

## Contributions

A.B. and P.T. designed the study; A.B. performed experiments, wrote code, and analyzed data; G.S. designed and performed the HCR *in situ* protocol; H.A. selected and provided the animals used; A.B., P.T., C.M., and C.Z. discussed the data and the analysis; C.M. suggested the smectic analogy and P.T. the Hydra embryo analogy. A.B. and P.T. wrote the manuscript, and all authors made comments.

## Acknowledgments

We thank Bert Hobmayer for generously guiding us into the Hydra world, helping with protocols, providing the lines used, and discussions, Taylor Skokan and Fabian Braukmann for help with the aggregate protocol, the Tomancak lab, Charisios Tsiairis, and Sera Weevers for discussions, Charlène Brillard for designing the cooler used during experiments, the MPI-CBG Light Microscopy Facility, in particular Jan Peychl, Britta Schroth-Diez and Riccardo Maraspini, for support, the MPI-CBG Fish facility for providing *Artemia nauplii* feeding Hydra, and Pierre Mangeol for critical reading of the manuscript. A.B. was supported by an EMBO LTF fellowship ALTF 785-2020, an ELBE postdoctoral fellowship, and MPI-CBG core funding.

---

**Fig. S1.**
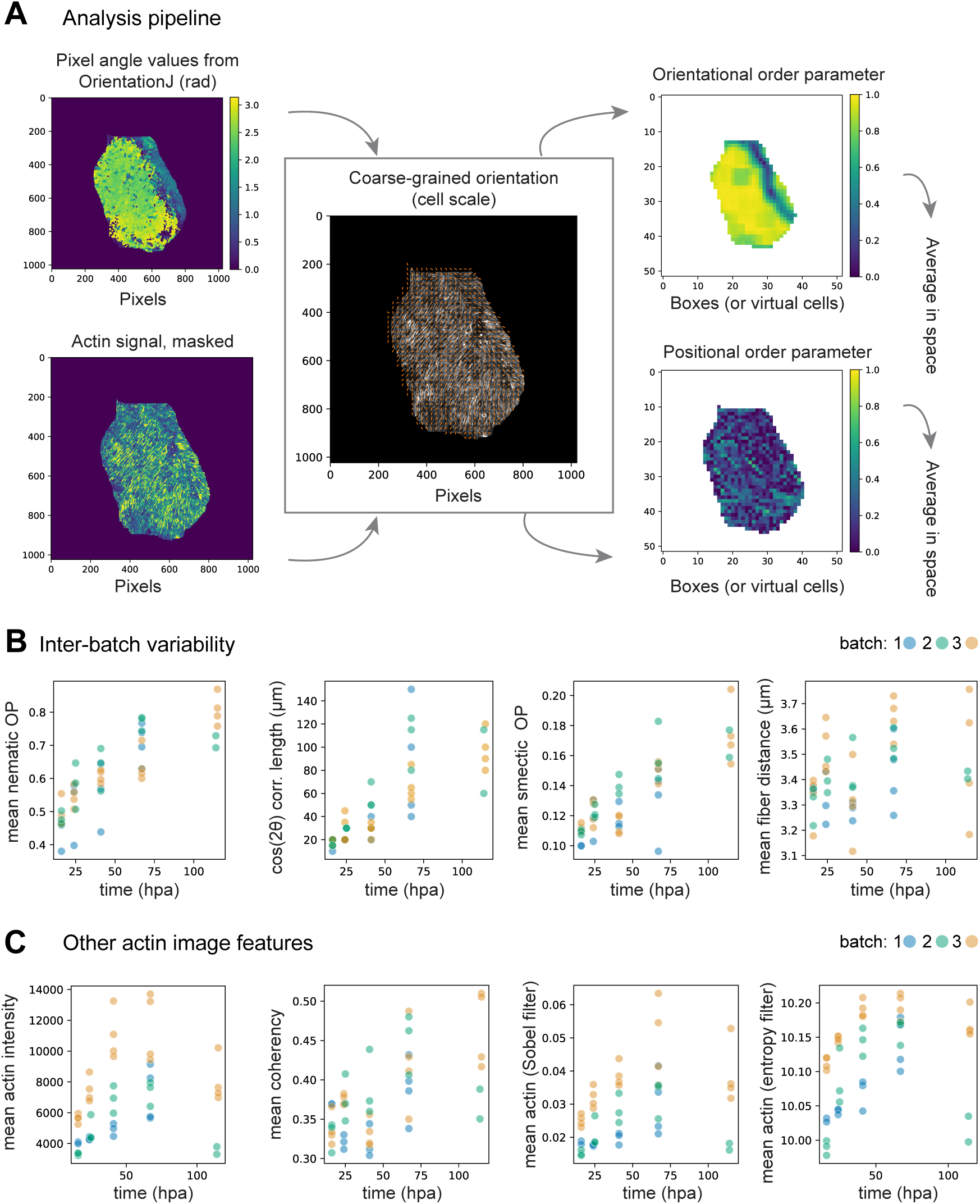
(**A**) Analysis pipeline principle. A map of the angle’s values at the fiber (i.e. 2-pixels) scale is obtained with OrientationJ. This map and a mask of the sample based on the actin signal are combined and coarse-grained at the cellular scale (central image). From this, the orientational and positional order parameter maps are obtained, and then averaged spatially. (**B**) The same plots as **Fig. 1C** and **D**, but with the batch of each aggregate color-coded. (**C**) Actin mean intensity, mean coherency, mean contrast (as measured with a Sobel filter and with an entropy filter), as a function of time. These measurements show different time evolution and differences between batches than the order parameters measurements.

**Fig. S2.**
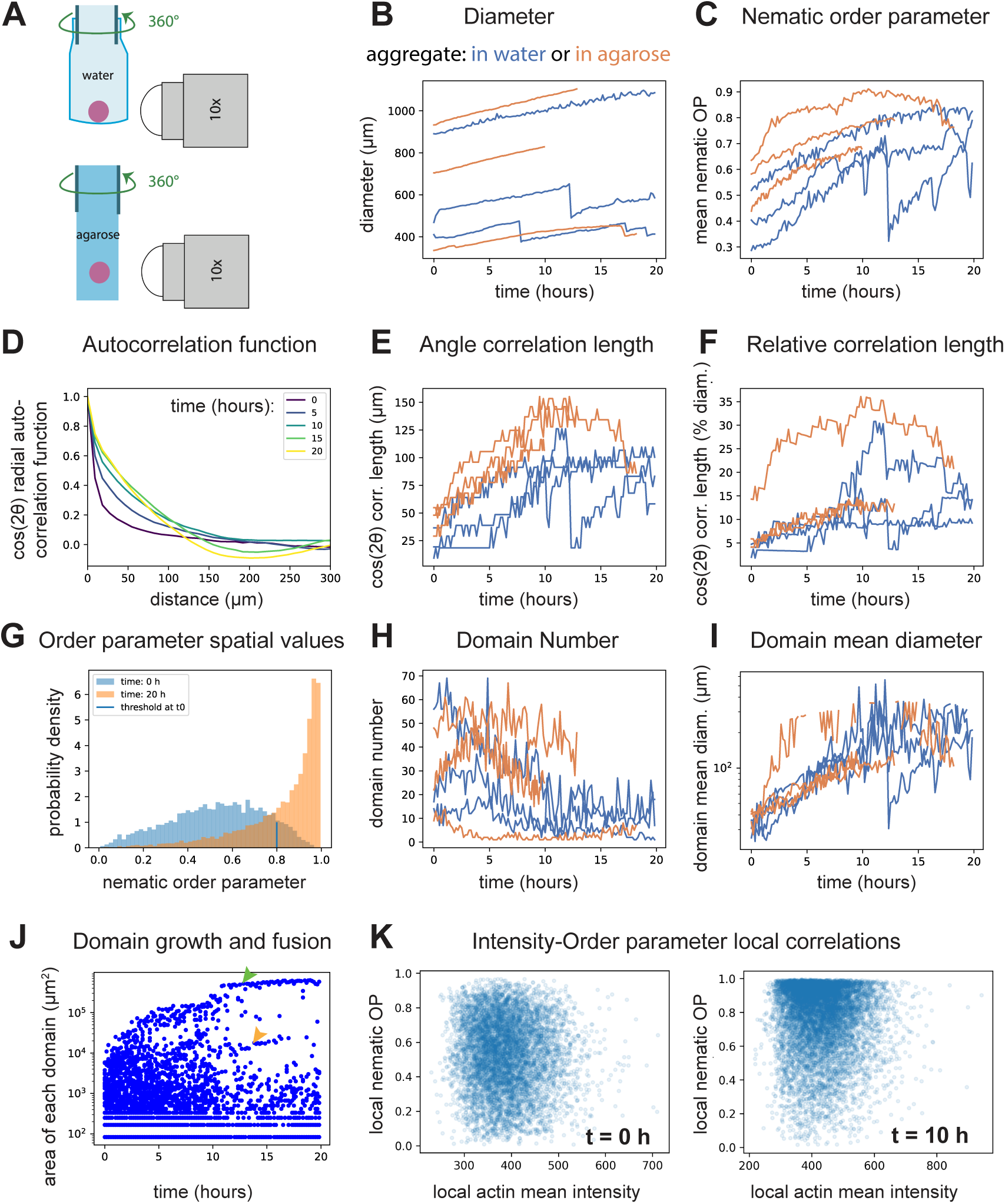
(**A**) Schematic of light-sheet sample mounting in water or agarose. The pink disks are aggregates, the green lines indicate glass capillary sections, and on the right is the detection objective. (**B**) Time evolution of the diameters of 3 live LifeAct-GFP aggregates in water (blue lines) and 3 in agarose (orange lines) from N=6 batches. (**C**) Spatial average of the orientational order parameter (nematic OP) with time of the same samples. (**D**) Autocorrelation function of cos(2θ) used to determine the correlation length in Fig. 2D, at 5 time points (in hpa) from the aggregate in Fig. 2A. (**E**) Correlation length (defined as in Fig. 1C) with time of the same 6 samples. (**F**) Correlation length normalized by the sample’s diameter at each time point. (**G**) Distribution of the spatial nematic order parameter values of the aggregate in Fig. 2A, with the 10% threshold used in the definition of the high-order domains (t0 is t=0h). After 20h, many more domains have reached the threshold. (**H**) High-order domains number with time of the same 6 samples. (**I**) Mean high-order domain diameters with time of the same 6 samples. (**J**) Individual areas of high-order domains with time in the aggregate in Fig. 2A. One dot is one domain at one time point. The arrows point at two identified domain trajectories, one of a big and one of a medium domain. (**K**) Correlations (or absence thereof) between actin intensity and order parameter at 0h (left) and 10h (right). One dot is one box (spatial area).

**Fig. S3.**
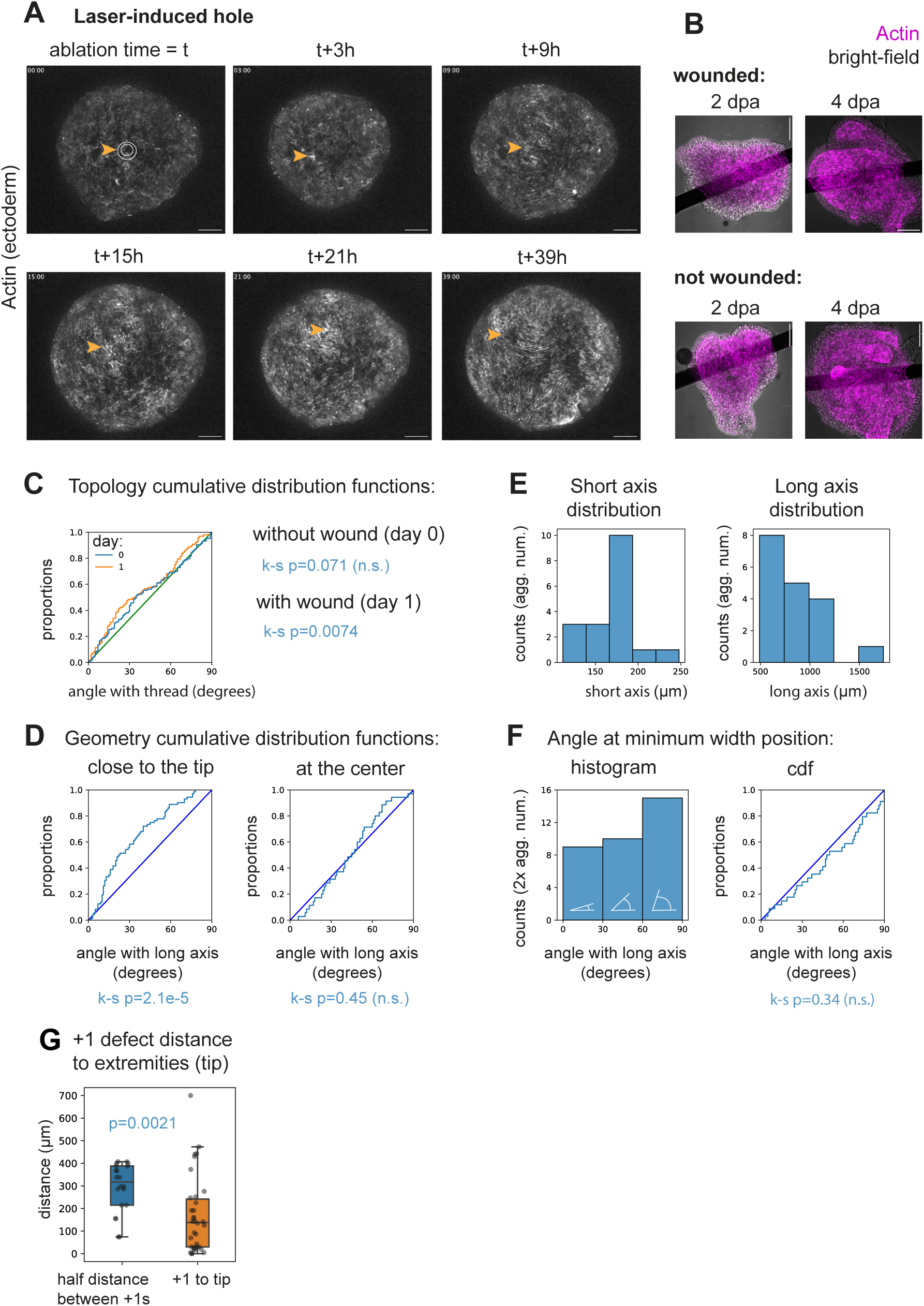
(**A**) Six time points of a spinning-disk microscopy movie of actin in the ectoderm (LifeAct-GFP) of an aggregate in agarose punctured with a laser at t=21 hpa. The orange arrows track the area of the puncture, circled in white on the first image (inner circle: initial puncture, outer circle: recoil). (**B**) Composite images of the bright field channel (black and white) and actin (magenta) of the aggregates in Fig. 3A,**B**. (**C**) Cumulative distribution functions (cdf) of all measurements used to produce the histograms in Fig. 3B (orange line) and d (blue line), and of the cdf of a uniform distribution (green line). (**D**) Cdf of all measurements used to produce the histograms in Fig. 3F and of the cdf of a uniform distribution (blue line). (**E**) Histograms of the short and long axes of the samples analyzed. (**F**) Manual measurements of the angle of actin with respect to the long axis at the position of minimum width along the same axis. Left: histogram, right: cdf. (**G**) Boxplot of half distance between +1 defects (left) and of distance between +1 and tip (right). One dot is one measurement point. All microscopy images are maximum-intensity stack projections. All scalebars are 100 microns.

**Fig. S4.**
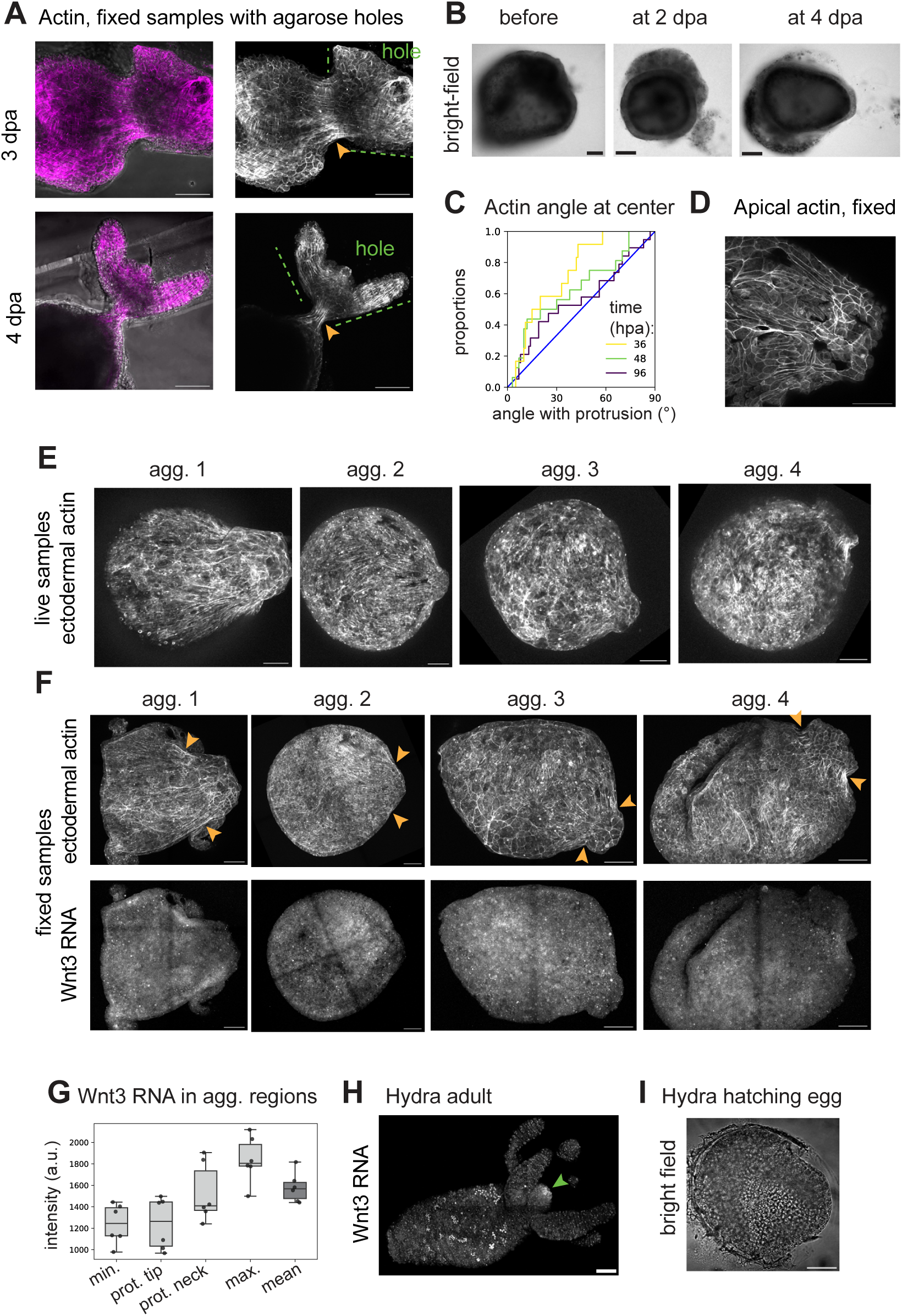
(**A**) Microscopy images of actin (phalloidin) in WT aggregates placed in agarose boxes with holes at 1 dpa and fixed at 3-4 dpa. Both examples show a striking alignment of actin at the neck (orange arrows). (**B**) Bright-field images of the aggregates shown in Fig. 4B**-D**. (**C**) Cumulative distribution function of the measurements used to produce the histograms in Fig. 4E. (**D**) Max. projection of the 4 most apical planes of protrusion 1 shown in Fig. 4G. (**E**) Live microscopy images of actin (LifeAct-GFP) in 4 of the aggregates of Fig. 4G before fixation. (**F**) Large-field images of these 4 aggregates. Top: actin (LifeActGFP). Orange arrows: neck sides. The protrusions can be identified with those in **e**. Bottom: raw Wnt3 RNA signal. (**G**) Boxplots of Fig. 4H before the normalization by the mean aggregate signal. (**H**) HCR RNA *in situ* detection of *Wnt3* on an adult Hydra. Green arrow: head organizer. The bright cells in the body and tentacles are auto-fluorescent nematoblasts. **i**. Bright-field image of the embryo in Fig. 4H. All fluorescence microscopy images are maximum-intensity stack projections. All scalebars are 100 microns.

## Movie Legends

**Movie S1.**

Example of a WT aggregate regeneration in water over days, bright-field imaging. Scalebar: 100 microns. Timestamps: hours after aggregation (hpa).

**Movie S2.**

Actin-tagged (LifeAct-GFP) aggregate regeneration in water from 28 hpa to 48 hpa, mounted in an FEP cuvette and imaged with a light-sheet microscope. Scalebar: 100 microns. Timestamps: hour:min, relative to the movie start.

**Movie S3.**

Nematic order parameter value calculated from the aggregate in Movie S2, superimposed on the actin Movie S2. X and Y axis: position in pixels. Colormap: nematic order parameter value.

**Movie S4.**

LifeAct-GFP aggregate regeneration after laser puncture, from 21 hpa to 63 hpa, mounted in 1% agarose and imaged with a spinning-disk microscope. Scalebar: 100 microns. Timestamps: hour:min, relative to the movie start.

**Movie S5.**

LifeAct-GFP aggregate regeneration in a 2% agarose box with a hole, imaged just after hole creation, from 22 hpa to 45 hpa, with a spinning-disk microscope. Scalebar: 100 microns. Timestamps: hour:min, relative to the movie start.

**Movie S6.**

Later LifeAct-GFP aggregates regeneration in an agarose box with a hole (after being imaged as in Movie S5), from 45 hpa to 88 hpa, bright-field imaging. Scalebar: 100 microns. Timestamps: hour:min, relative to the movie start. The aggregates can move inside the agarose (left aggregate), or stay relatively still (right aggregate).

## Methods

### Hydra culture

*Hydra vulgaris* lines are cultured according to standard protocols (Seybold et al., 2016) at 17± 1°C in Hydra Medium, HM (1mM Tris, 1mM CaCl2, 0.4mM MgCl2, 0.4mM KCL, 4mM NaHCO3, pH=7.3-7.6) and fed once or twice a week with artemia nauplii less than 48 hours old. The lines used are the *Hydra magnipapillata* 105 as WT and the *Hydra vulgaris* actin1-Lifeact-EGFP line inserted in the ectoderm (Aufschnaiter et al., 2017), both gifts from the Hobmayer lab.

### Aggregates preparation

To prepare aggregates, 50 Hydras fed 48 to 72 hours earlier are selected. The body column parts are dissected by cutting below the tentacles and above the bud zone. The following steps are performed at 18°C using a custom-made Peltier tube cooler. The body columns are incubated in Dissociation Medium, DM (3.6 mM KCL, 6 mM CaCl2, 1.2 mM MgSO4, 6 mM NaCitrat, 6 mM NaPyruvat, 4 mM Glucose, 12.5 mM TES, 50 mg/L Kanamycin 100 mg/L Streptomycin, pH 6.9) during 20 min. The DM is removed, 1.5 mL of new DM is added, and then the body columns are gently pipetted up and down with a flamed glass pipette for 30 sec, the solution is left to sediment for 2 min, and the supernatant containing single cells and small cell clusters is collected. The cycle of the pipetting-supernatant collection is repeated around 8 times until the pieces of tissue are fully dissociated. The resulting cell suspension is centrifuged at 150 g during 5 min at 4°C, the supernatant discarded, and the cell pellet resuspended in 4mL DM, aliquoted in Eppendorf tubes, and centrifuged again at 150 g during 5 min at 4°C. The tubes are placed face down in a Petri dish containing DM until the cell pellets detach and sink into the Petri dish. The pellets are cut in 2, with *in fine* in a ratio of 2-3 Hydra body columns used per aggregate. After 2 hours, the aggregates are transferred into a 1:1 HM-DM mix, in which they stay overnight. The next day, they are transferred into a 3:1 HM-DM mix, then around 6 hours later into pure HM. For 2 out of 3 samples used in Fig. 2, full Hydras were used instead of body columns, and the cell suspension was filtered using a 100 microns diameter sieve before the first centrifugation, making sure to remove big clusters (epithelial cells are 20 to 30 microns in diameter), and the aggregate development appeared similar.

### Perturbations of aggregates topology, geometry, and deformation

#### ‘skewer’ aggregates (Fig. 3)

The aggregates are made as described and pierced using a 50-micron diameter nylon thread with the help of tweezers, either immediately after being made or the day later. Changes of medium proceed normally, but the medium is removed instead of the aggregates being transferred to a new dish.

#### ‘sausage’ aggregates (Fig. 3)

The cell suspension is prepared as described until the end of the first centrifugation. The supernatant is then discarded, but no DM is added. The cell suspension is transferred to 200 microns inner diameter glass capillaries, and the capillaries are plugged with dentist paste and then centrifuged at 150 g for 5 min at 4°C. The capillaries are kept for one hour at 18°C, and then the elongated pellets (‘sausages’) are pushed out of the capillaries into a DM-filled Petri dish. After 1 hour, the ‘sausages’ are transferred into 1:1 HM-DM. 2 hours later, they are embedded into 2% low-melting agarose drops, covered with 1:1 HM-DM, where they stay overnight. The next day, the solution is replaced with 3:1 HM-DM. At around 20 hpa, the agarose shell is broken using needles, and the freed aggregates are transferred into pure HM.

#### ‘boxed’ aggregates (Fig. 4)

The aggregates are prepared as described. The next day at around 24 hpa, they are embedded in 1 or 2% low-melting agarose, in direct contact with the tip of a 200 microns diameter nylon thread. After the agarose solidifies, the thread is then gently pulled away, exerting some force on the aggregate and leaving a cylindrical hole of 200 microns in diameter into the agarose surrounding the aggregate.

### Fixation and staining

#### Fixation

The aggregates are fixed using fresh 4% PFA in HM at room temperature for 45 min. They are rinsed twice in HM, washed for 5 min in HM, washed for 5 min in PBS, and stored at 4 degrees in PBS. For HCR *in situs*, the PBS wash is replaced by a 1:1 MeOH-HM 5min wash and a 100% MeOH 5min wash, and the samples are stored at −20°C in MeOH.

#### Phalloidin staining

After fixation, the samples are incubated for 1h30 in PBST (PBS + 0.05% Triton X-100) at room temperature. Then, they are incubated in PBST with DAPI (1:250) and rhodamine-phalloidin (1:250) at 4°C overnight on a rocking platform. The next day, they are rinsed twice with PBS, washed 2×5 min in PBS, soaked in glycerol (80%)-DABCO overnight, mounted in between two coverslips with tape as spacers, and imaged within a few days from both sides.

### Whole-mount HCR RNA *in situ* hybridization

Hybridization Chain Reaction RNA *in situ* against *HyWnt3* was performed according to published protocols (Choi et al., 2018; Vellutini et al., 2023) with the following modifications: the protocol always started from fixed aggregates conserved in 100% MeOH at −20C for at least 2-3 days. After the rehydration steps, the samples were incubated for 5 min at room temperature in 1 mL of 10 μg/mL Protease K solution in PBS-Tween 20 (PBT) 0.1%. After the post-fixation step in 4% PFA in PBT 0.1%, the samples were washed 5 times for 5 min in 1 mL PBT 0.1% and once for 5 min in 500 μL 2x SSC-Tween 20 0.1%. The pre-hybridization took place in a volume of 400 μL of probe hybridization buffer, while the O/N incubation with the RNA probes took place in a total volume of 200 μL.

### Fixed imaging

Fluorescently labeled fixed samples (Fig. 1,3, and 4G) were imaged using a Zeiss LSM 700 microscope equipped with a Zeiss Axio Observer.Z1 inverted stand and a 25x (NA=0.8) multi-immersion objective in glycerol or water depending on the sample, and operated with Zeiss ZEN 2012 SP5 FP3 (black). *LifeAct GFP (Fig. 1):* LifeAct-GFP fixed aggregates were imaged with a 488nm DPSS laser, a zoom factor of 0.5x, a pixel size of 0.5 microns, and a stack covering from the surface until the equator of the aggregate (around 50 to 100 microns thick), with a z plane spacing of 1.49 microns. Glycerol solution was used for 2 of the batches and water for one of the batches, without any strong visible difference. *Phalloidin+BF+Dapi (Fig. 3):* WT fixed aggregates were imaged in glycerol using a laser diode 405 nm (for DAPI), a laser DPSS 555 nm (for rhodamin phalloidin), and a laser DPSS 639 nm (for bright-field), a zoom factor of 0.5x, a pixel size of 0.5 to 1.0 microns and a stack covering from the surface until the equator of the aggregate (around 50 to 100 microns thick), with a z-plane spacing of 1.31 microns. *HCR in situs (Fig. 4):* LifeAct-GFP fixed aggregates were imaged in glycerol with a 488nm DPSS laser (for LifeAct-GFP), a laser DPSS 639 nm (for HyWnt3-Alexa647nm). For large-scale views, a zoom factor of 0.5x, a pixel size of 0.57 microns, and a stack covering from the surface until the equator of the aggregate (around 40 to 80 microns thick), with a z-plane spacing of 2.0 microns, were used. For zoomed-in views, a zoom factor of 0.6 to 1x, a pixel size of 0.15 to 0.21 microns, and a z spacing of 1.49 microns were used.

### Live imaging

#### SPIM (Fig. 2)

samples were imaged using a Zeiss Light-sheet Z.1 with Zeiss 10x (0.2 NA) Air illumination objectives and a Zeiss Plan Apochromat 10x (0.5 NA) water immersion detection objective, operated with Zeiss ZEN 2014 SP1. Samples free-floating in HM were mounted in custom cubic cuvettes made of 0.05mm-thick FEP foil, with a ∼1.5×1.5 mm^2^ section, and attached to a glass capillary. Samples in agarose were mounted using glass capillaries (2 mm outer diameter, ∼1.5mm inner) filled with 1% low-melting agarose according to standard protocols. Both were imaged using a 0.5-1x zoom factor according to the sample size (which results in a pixel size from 0.46 to 0.91 microns), filling up the chamber with Hydra Medium (HM) kept at 18°C. Four angles view stacks were acquired by exciting the EGFP with a 488nm DPSS laser at low power, with 30ms acquisition time per frame and one plane every 6 microns covering around 540 microns (depending on the sample size). One time point per 10 min was acquired to minimize laser exposure of the sample.

#### Spinning disk (Fig. S2, Fig. 4)

samples were imaged using a Zeiss Spinning Disk equipped with a Yokogawa CSU-X1 scan head (5000 rpm, pinhole diameter 50 μm, pinhole spacing 250 μm) and a Zeiss AxioCam 705 Mono CCD camera, and a Zeiss Plan Apochromat 10x (0.45 NA) Air objective. The samples were mounted in a glass-bottom MatTek 35mm dish, covered with 1 or 2% agarose and HM, and kept at 18°C using a Warner cooling chamber. LifeAct-EGFP was excited with a laser DPSS 488 nm 200ms per frame. A pixel size of 0.69 microns and a z-step of 3.0 microns were used, covering around 300 microns in depth. One time point per 10 min was acquired, in order to minimize laser exposure of the sample.

#### Laser ablation

In combination with the spinning disk system described above, RAPP optoelectronic 355nm laser (1 kHz rep rate, max 42 μJ pulse power, 1 ns pulse length) operated by Syscon2 was used to cut out a disk of diameter in the epithelium. The disk cutting itself lasts ∼100-150ms in total, and we stream recorded (as fast as possible) during the ablation, with a 100ms exposure time. Long movies were then acquired as above with a 10-minute time step.

### Image analysis pipeline

The goal of this pipeline is to extract the coarse-grained actin orientation and measure order parameters, correlation lengths, and high-order domain properties from maximum intensity projections of the stacks of actin signal. The initial angle measurement is done using Fiji and OrientationJ (Püspöki et al., 2016), while the coarse-graining and further measurements are made with custom Python codes. For fixed images, both sides of the aggregates were analyzed and averaged (Fig. 1). For live imaging (Fig. 2), one side of each aggregate was chosen.

#### Angles measurement

The OrientationJ ‘OrientationJAnalysis’ plugin (Püspöki et al., 2016) was used to obtain the dominant angle of the image features. Briefly, this plugin calculates the image structure tensor from the gradient of the intensity within a window and its eigenvectors to obtain the Coherency and Angle, indicating resp. the magnitude and principal direction of the image gradient. A window with σ = 2 pixels was used to match the typical width of the actin fibers.

#### Mask of the aggregate (region of interest)

For live samples (Fig. 2), a mask is created in Fiji using a Triangle threshold segmentation of the actin movie, followed by erosion, dilation, and filling hole binary operations. For fixed samples (Fig. 1), the mask is created using Python, first applying a Gaussian blur of sigma=3 to the actin image, then segmenting the blurred image with a threshold of 200 (chosen after manual inspection of the mask obtained for the different samples), then performing 10 iterations of an erosion binary operation done with 20×20 matrices of ones element. The goal of this erosion is to remove the edges of the sample, which do not show the basal actin fibers.

#### Coarse-graining of angles values

To extract the average actin fibers angle at a cellular (rather than at a single fiber-) scale, we coarse-grained the Angle matrix obtained by OrientationJ in boxes of 10×10 microns^2^. We implemented a nematic version in Python by calculating the local director angle of each box according to 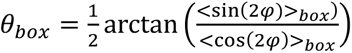, where *ϕ* is the angle measured at a 2-pixels scale by OrientationJ, the average is performed over the box region, and the numpy.arctan2 function was used.

#### Actin signal measurements

Actin intensities are coarse-grained by simply downscaling the raw image, and the average of the average in the mask area is calculated.

#### Correlation length

From the coarse-grained angle field θ(x,y), and since we are interested in the nematic properties, we calculated C(x,y), the autocorrelation matrix of cos(2θ) using the ITK library in Python and its method MaskedFFTNormalizedCorrelationImageFilter. Note that this method performs the autocorrelation on the masked image, which is important for the correlation length to fall to zero at a length larger than the object size. We then converted it to polar coordinate as C(r, θ) and averaged it radially (i.e., over θ) to obtain the autocorrelation function F(r), with r the distance between points. The correlation length was extracted by either fitting a decreasing exponential to the initial portion of the curve (over 6 microns), as the curve is only exponentially decreasing initially (Fig. S2d) or setting a threshold of 0.2 (i.e., 20% correlation) as the value of correlation corresponding to the characteristic correlation length. Both gave qualitatively similar results (increasing with time), but we here report the threshold of 0.2 length, which reflects better the non-zero correlation remaining at larger distances.

#### Nematic order parameter

The nematic order parameter was calculated as *S* =< cos (2(*θ* − *θ̂*) >*_neighb_*, where *θ̂* is the neighborhood director calculated as 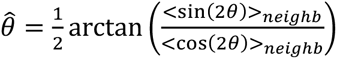. The neighborhood is chosen to be 50×50 microns^2^, corresponding to a cell and its immediate neighbors.

#### Smectic order parameter

The smectic order parameter would, in principle, be the amplitude of the principal mode of a Fourier decomposition of the actin density. To adapt it to our biological data, we devised a related smectic order parameter, easier to measure, as follows. For each box of 10×10 microns^2^, we use the corresponding angle value to select a 10 microns x 50 microns image stripe with its long axis orthogonal to the actin fibers. This becomes a 1D intensity profile I(r) by integrating over the smaller dimension (the one parallel to the nematic orientation). We derivate it to remove contributions from large-scale intensity variations, then calculate dI/dr autocorrelation function with numpy.correlate, normalize it to 1, and the height of the first autocorrelation peak after the r=0 peak is defined as the order parameter, while its position gives the characteristic spacing between the fibers.

#### High-order domains analysis

The threshold for ‘High-order domain’ is defined by the 10% of highest nematic order parameter values cut-off at time 0. The order parameter images at different time points are thresholded accordingly and the connected domains are identified using skimage.measure.label. This provides the number of domains with time. Their area is measured by counting the number of pixels of a given connected domain.

#### Homogeneous growth scenario

To provide the equivalent hypothetical homogeneous increase scenario, the spatially averaged order parameter time evolution is smoothed with a uniform filter of size 5, and differentiated to obtain the average rate of increase 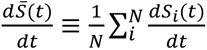 (where the sum is over the boxes *i*). In our homogeneous increase scenario, for a given time *t* we chose all the boxes to have the same rate of increase 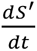 except the *n*_1_ boxes which have already reached 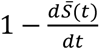, such that 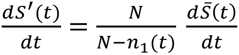. The order parameter evolution for the boxes is obtained iteratively with:

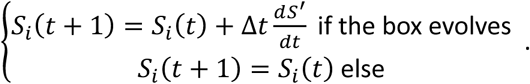

The synthetic images obtained are then analyzed as described above, and the area values are corrected by multiplying by *A*(*t*)⁄*A*(*t* = 0), where A is the total area of the sample, to take into account the effect of aggregate inflation.

### Manual analysis of actin angle

For perturbation conditions, the ectoderm actin orientation is measured manually within some ROIs using the ‘angle’ tool from Fiji.

#### ‘skewer’ aggregates (Fig. 3)

The ROIs are chosen to be well-aligned actin zones in proximity (∼50 microns in the direction of the other thread tip) to where the nylon thread enters the aggregate (as seen in bright-field images). For one aggregate, 2 such ROIs per side are measured, resulting in 4 measurements per aggregate. The angle is measured as the angle with the nylon thread, 0° being parallel to it, and 90° orthogonal to it.

#### ‘sausage’ aggregates (Fig. 3)

As the sausages are often bent at 4 dpa, the longest curvilinear axis is first defined manually using the ‘segmented line’ Fiji tool, and its length is measured. The ROIs angles are measured with 0° being parallel to the longest curvilinear axis and 90° orthogonal to it. A total of four measurements are made at the tips of each aggregate (2 sides x 2 tips). The midpoint ROI is estimated manually as the midway between the 2 tips along the curvilinear axis. Following this procedure, a total of two measurements is acquired per each aggregate (1 on each side). The orthogonal axis length is defined as the average between the shortest orthogonal (to the long axis) dimension and the largest one, as the sausages are not always of constant width.

#### ‘boxed’ aggregates (Fig. 4)

The central ROI is defined as the region between the center of the aggregate and ∼1/3^rd^ of the center-to-protrusion distance. Angles are measured with 0° being parallel to the center-to-protrusion line and 90° orthogonal to it. The neck ROI is defined as the beginning of the protrusion, where the cells are the thinnest. For later aggregates (2 and 4 dpa), the neck region is only defined if a protrusion is present, which is rare.

### Manual analysis of Wnt intensity

Image stacks are stitched using the Zen software then a maximum intensity projection is performed. The HCR *in situ* signal of *Wnt3* is characterized by small bright dots. Thus, to remove the background signal, the *Wnt* maximum projection images are processed using a 20-pixel rolling ball background subtraction (using Fiji). The ROI for the protrusion tip and neck are defined on the actin channel, with the tip being the region with rounded cells and the neck with elongated actin fibers. The average intensity is then measured using the same ROIs on the *Wnt* channel with the Fiji measurement tool. For comparison, two large ROIs are defined elsewhere in the actin channel, and their respective average actin intensity is measured on the *Wnt* channel. The maximum *Wnt* intensity ROI is chosen manually on the *Wnt* channel to be the region with seemingly the most actin and diameter similar to that of the neck ROI. The minimum *Wnt* intensity ROI is defined similarly but on the seemingly minimum signal with a diameter similar to the protrusion region. All ROIs are chosen to avoid the stitching area artifact (cross-shaped lower-intensity regions generated by the image stitching process).

### Statistics

To assess the randomness of actin orientation upon perturbations (Fig. 3 and Fig. 4), the measurement distribution was normalized in the range 0-1 and tested against a uniform distribution with a Kolmogorov-Smirnov test using the Python function scipy.stats.kstest. All the measurement points per sample were used assuming they are independent, as they are >200 microns from each other, the distance over which correlation in angle orientation disappears (Fig. 1). To assess the difference between the distributions in Fig. 3H, we performed a Mann-Whitney test using the function scipy.stats.mannwhitneyu.

## Bibliography

Aufschnaiter, R., Wedlich-Söldner, R., Zhang, X., Hobmayer, B., 2017. Apical and basal epitheliomuscular F-actin dynamics during *Hydra* bud evagination. Biology Open bio.022723. 10.1242/bio.022723

Bade, N.D., Kamien, R.D., Assoian, R.K., Stebe, K.J., 2017. Curvature and Rho activation differentially control the alignment of cells and stress fibers. Science Advances 3, e1700150. 10.1126/sciadv.1700150

Bode, H., 2011. Axis Formation in Hydra. Annu. Rev. Genet. 45, 105–117. 10.1146/annurev-genet-102209-163540

Braun, E., Keren, K., 2018. *Hydra* Regeneration: Closing the Loop with Mechanical Processes in Morphogenesis. BioEssays 40, 1700204. 10.1002/bies.201700204

Broun, M., Gee, L., Reinhardt, B., Bode, H.R., 2005. Formation of the head organizer in hydra involves the canonical Wnt pathway. Development 132, 2907–2916. 10.1242/dev.01848

Burridge, K., Guilluy, C., 2016. Focal adhesions, stress fibers and mechanical tension. Experimental Cell Research 343, 14–20. 10.1016/j.yexcr.2015.10.029

Cazet, J.F., Cho, A., Juliano, C.E., 2021. Generic injuries are sufficient to induce ectopic Wnt organizers in Hydra. eLife 10, e60562. 10.7554/eLife.60562

Choi, H.M.T., Schwarzkopf, M., Fornace, M.E., Acharya, A., Artavanis, G., Stegmaier, J., Cunha, A., Pierce, N.A., 2018. Third-generation in situ hybridization chain reaction: multiplexed, quantitative, sensitive, versatile, robust. Development 145, dev165753. 10.1242/dev.165753

Clevers, H., 2016. Modeling Development and Disease with Organoids. Cell 165, 1586–1597. 10.1016/j.cell.2016.05.082

Cochet-Escartin, O., Locke, T.T., Shi, W.H., Steele, R.E., Collins, E.-M.S., 2017. Physical Mechanisms Driving Cell Sorting in Hydra. Biophysical Journal 113, 2827–2841. 10.1016/j.bpj.2017.10.045

Collinet, C., Lecuit, T., 2021. Programmed and self-organized flow of information during morphogenesis. Nat Rev Mol Cell Biol 22, 245–265. 10.1038/s41580-020-00318-6

del Rio, A., Perez-Jimenez, R., Liu, R., Roca-Cusachs, P., Fernandez, J.M., Sheetz, M.P., 2009. Stretching Single Talin Rod Molecules Activates Vinculin Binding. Science 323, 638–641. 10.1126/science.1162912

Elsdale, T., Wasoff, F., 1976. Fibroblast cultures and dermatoglyphics: The topology of two planar patterns. Wilhelm Roux’ Archiv 180, 121–147. 10.1007/BF00848102

Fenix, A.M., Neininger, A.C., Taneja, N., Hyde, K., Visetsouk, M.R., Garde, R.J., Liu, B., Nixon, B.R., Manalo, A.E., Becker, J.R., Crawley, S.W., Bader, D.M., Tyska, M.J., Liu, Q., Gutzman, J.H., Burnette, D.T., 2018. Muscle-specific stress fibers give rise to sarcomeres in cardiomyocytes. eLife 7, e42144. 10.7554/eLife.42144

Ferenc, J., Papasaikas, P., Ferralli, J., Nakamura, Y., Smallwood, S., Tsiairis, C.D., 2021. Mechanical oscillations orchestrate axial patterning through Wnt activation in *Hydra*. Sci. Adv. 7, eabj6897. 10.1126/sciadv.abj6897

Fütterer, C., Colombo, C., Jülicher, F., Ott, A., 2003. Morphogenetic oscillations during symmetry breaking of regenerating *Hydra vulgaris* cells. Europhys. Lett. 64, 137–143. 10.1209/epl/i2003-00148-y

Galliot, B., 2012. Hydra, a fruitful model system for 270 years. Int. J. Dev. Biol. 56, 411–423. 10.1387/ijdb.120086bg

Gee, L., Hartig, J., Law, L., Wittlieb, J., Khalturin, K., Bosch, T.C.G., Bode, H.R., 2010. β-catenin plays a central role in setting up the head organizer in hydra. Developmental Biology 340, 116–124. 10.1016/j.ydbio.2009.12.036

Gierer, A., Berking, S., Bode, H., David, C.N., Flick, K., Hansmann, G., Schaller, H., Trenkner, E., 1972. Regeneration of Hydra from Reaggregated Cells. Nature New Biology 239, 98–101. 10.1038/newbio239098a0

Greenberg, M.J., Arpağ, G., Tüzel, E., Ostap, E.M., 2016. A Perspective on the Role of Myosins as Mechanosensors. Biophysical Journal 110, 2568–2576. 10.1016/j.bpj.2016.05.021

Guillamat, P., Blanch-Mercader, C., Pernollet, G., Kruse, K., Roux, A., 2022. Integer topological defects organize stresses driving tissue morphogenesis. Nat. Mater. 21, 588–597. 10.1038/s41563-022-01194-5

Han, M.K.L., Rooij, J. de, 2016. Converging and Unique Mechanisms of Mechanotransduction at Adhesion Sites. Trends in Cell Biology 26, 612–623. 10.1016/j.tcb.2016.03.005

Harris, A.R., Jreij, P., Fletcher, D.A., 2018. Mechanotransduction by the Actin Cytoskeleton: Converting Mechanical Stimuli into Biochemical Signals. Annual Review of Biophysics 47, 617–631. 10.1146/annurev-biophys-070816-033547

Hobmayer, B., Rentzsch, F., Kuhn, K., Happel, C.M., Von Laue, C.C., Snyder, P., Rothbächer, U., Holstein, T.W., 2000. WNT signalling molecules act in axis formation in the diploblastic metazoan Hydra. Nature 407, 186–189. 10.1038/35025063

Huycke, T.R., Miller, B.M., Gill, H.K., Nerurkar, N.L., Sprinzak, D., Mahadevan, L., Tabin, C.J., 2019. Genetic and Mechanical Regulation of Intestinal Smooth Muscle Development. Cell 179, 90–105.e21. 10.1016/j.cell.2019.08.041

Kücken, M., Soriano, J., Pullarkat, P.A., Ott, A., Nicola, E.M., 2008. An Osmoregulatory Basis for Shape Oscillations in Regenerating Hydra. Biophysical Journal 95, 978–985. 10.1529/biophysj.107.117655

Livne, A., Geiger, B., 2016. The inner workings of stress fibers − from contractile machinery to focal adhesions and back. Journal of Cell Science 129, 1293–1304. 10.1242/jcs.180927

Livshits, A., Shani-Zerbib, L., Maroudas-Sacks, Y., Braun, E., Keren, K., 2017. Structural Inheritance of the Actin Cytoskeletal Organization Determines the Body Axis in Regenerating Hydra. Cell Reports 18, 1410–1421. 10.1016/j.celrep.2017.01.036

López-Gay, J.M., Nunley, H., Spencer, M., di Pietro, F., Guirao, B., Bosveld, F., Markova, O., Gaugue, I., Pelletier, S., Lubensky, D.K., Bellaïche, Y., 2020. Apical stress fibers enable a scaling between cell mechanical response and area in epithelial tissue. Science 370, eabb2169. 10.1126/science.abb2169

Mao, Q., Acharya, A., Rodríguez-delaRosa, A., Marchiano, F., Dehapiot, B., Al Tanoury, Z., Rao, J., Díaz-Cuadros, M., Mansur, A., Wagner, E., Chardes, C., Gupta, V., Lenne, P.-F., Habermann, B.H., Theodoly, O., Pourquié, O., Schnorrer, F., 2022. Tension-driven multi-scale self-organisation in human iPSC-derived muscle fibers. eLife 11, e76649. 10.7554/eLife.76649

Maroudas-Sacks, Y., Garion, L., Shani-Zerbib, L., Livshits, A., Braun, E., Keren, K., 2021. Topological defects in the nematic order of actin fibres as organization centres of Hydra morphogenesis. Nat. Phys. 17, 251–259. 10.1038/s41567-020-01083-1

Maroudas-Sacks, Y., Garion, L., Suganthan, S., Popović, M., Keren, K., 2024a. Confinement Modulates Axial Patterning in Regenerating Hydra. PRX Life 2, 043007. 10.1103/PRXLife.2.043007

Maroudas-Sacks, Y., Suganthan, S., Garion, L., Ascoli-Abbina, Y., Westfried, A., Dori, N., Pasvinter, I., Popović, M., Keren, K., 2024b. Mechanical strain focusing at topological defect sites in regenerating Hydra. 10.1101/2024.06.13.598802

Martin, V.J., Littlefield, C.L., Archer, W.E., Bode, H.R., 1997. Embryogenesis in Hydra. The Biological Bulletin 192, 345–363. 10.2307/1542745

Mercker, M., Köthe, A., Marciniak-Czochra, A., 2015. Mechanochemical Symmetry Breaking in Hydra Aggregates. Biophysical Journal 108, 2396–2407. 10.1016/j.bpj.2015.03.033

Mombach, J.C.M., De Almeida, R.M.C., Thomas, G.L., Upadhyaya, A., Glazier, J.A., 2001. Bursts and cavity formation in Hydra cells aggregates: experiments and simulations. Physica A: Statistical Mechanics and its Applications 297, 495–508. 10.1016/S0378-4371(01)00199-6

Noda, K., 1970. The Fate of Aggregates formed by Two Species of Hydra (Hydra magnipapillata and Pelmatohydra robusta).

Pukhlyakova, E., Aman, A.J., Elsayad, K., Technau, U., 2018. β-Catenin–dependent mechanotransduction dates back to the common ancestor of Cnidaria and Bilateria. Proc. Natl. Acad. Sci. U.S.A. 115, 6231–6236. 10.1073/pnas.1713682115

Püspöki, Z., Storath, M., Sage, D., Unser, M., 2016. Transforms and Operators for Directional Bioimage Analysis: A Survey, in: De Vos, W.H., Munck, S., Timmermans, J.-P. (Eds.), Focus on Bio-Image Informatics. Springer International Publishing, Cham, pp. 69–93. 10.1007/978-3-319-28549-8_3

Ravichandran, Y., Vogg, M., Kruse, K., Pearce, D.J., Roux, A., 2024. Topology changes of the regenerating Hydra define actin nematic defects as mechanical organizers of morphogenesis.

Sato-Maeda, M., Tashiro, H., 1999. Development of Oriented Motion in Regenerating Hydra Cell Aggregates. Zoological Science 16, 327–334. 10.2108/zsj.16.327

Seybold, A., Salvenmoser, W., Hobmayer, B., 2016. Sequential development of apical-basal and planar polarities in aggregating epitheliomuscular cells of Hydra. Developmental Biology 412, 148–159. 10.1016/j.ydbio.2016.02.022

Shani-Zerbib, L., Garion, L., Maroudas-Sacks, Y., Braun, E., Keren, K., 2022. Canalized Morphogenesis Driven by Inherited Tissue Asymmetries in Hydra Regeneration. Genes 13, 360. 10.3390/genes13020360

Shyer, A.E., Rodrigues, A.R., Schroeder, G.G., Kassianidou, E., Kumar, S., Harland, R.M., 2017. Emergent cellular self-organization and mechanosensation initiate follicle pattern in the avian skin. Science 357, 811–815. 10.1126/science.aai7868

Skokan, T.D., Vale, R.D., McKinley, K.L., 2020. Cell Sorting in Hydra vulgaris Arises from Differing Capacities for Epithelialization between Cell Types. Current Biology 30, 3713–3723.e3. 10.1016/j.cub.2020.07.035

Sonam, S., Balasubramaniam, L., Lin, S.-Z., Ivan, Y.M.Y., Pi-Jaumà, I., Jebane, C., Karnat, M., Toyama, Y., Marcq, P., Prost, J., Mège, R.-M., Rupprecht, J.-F., Ladoux, B., 2023. Mechanical stress driven by rigidity sensing governs epithelial stability. Nat. Phys. 19, 132–141. 10.1038/s41567-022-01826-2

Soriano, J., Rüdiger, S., Pullarkat, P., Ott, A., 2009. Mechanogenetic Coupling of Hydra Symmetry Breaking and Driven Turing Instability Model. Biophysical Journal 96, 1649–1660. 10.1016/j.bpj.2008.09.062

Vellutini, B.C., Cuenca, M.B., Krishna, A., Szałapak, A., Modes, C.D., Tomančák, P., 2023. Patterned embryonic invagination evolved in response to mechanical instability. 10.1101/2023.03.30.534554

Vogg, M.C., Galliot, B., Tsiairis, C.D., 2019. Model systems for regeneration: *Hydra*. Development 146, dev177212. 10.1242/dev.177212

Wang, R., Steele, R.E., Collins, E.-M.S., 2020. Wnt signaling determines body axis polarity in regenerating Hydra tissue fragments. Developmental Biology 467, 88–94. 10.1016/j.ydbio.2020.08.012

Wang, Z., Marchetti, M.C., Brauns, F., 2023. Patterning of morphogenetic anisotropy fields. Proceedings of the National Academy of Sciences 120, e2220167120. 10.1073/pnas.2220167120

Weevers, S.L., Falconer, A.D., Mercker, M., Sadeghi, H., Ferenc, J., Ott, A., Oelz, D.B., Marciniak-Czochra, A., Tsiairis, C.D., 2024. Mechanochemical Patterning Localizes the Organizer of a Luminal Epithelium. 10.1101/2024.10.29.620841

Yang, Q., Liberali, P., 2021. Collective behaviours in organoids. Current Opinion in Cell Biology, Cell Dynamics 72, 81–90. 10.1016/j.ceb.2021.06.006

Yevick, H.G., Duclos, G., Bonnet, I., Silberzan, P., 2015. Architecture and migration of an epithelium on a cylindrical wire. Proceedings of the National Academy of Sciences 112, 5944–5949. 10.1073/pnas.1418857112

